# FliW and CsrA govern flagellin (FliC) synthesis and play pleiotropic roles in virulence and physiology of *Clostridioides difficile* R20291

**DOI:** 10.1101/2021.05.06.443043

**Authors:** Duolong Zhu, Shaohui Wang, Xingmin Sun

**Author notes:** Corresponding author. Address: 12901 Bruce B. Downs Blvd Tampa, FL 33612, United States.

## Abstract

*Clostridioides difficile* is a Gram-positive, spore-forming, and toxin-producing anaerobe that can cause nosocomial antibiotic-associated intestinal disease. In *C. difficile*, the expression of flagellar genes is coupled to toxin gene regulation and bacterial colonization and virulence. The flagellin FliC is responsible for pleiotropic gene regulation during *in vivo* infection. However, how *fliC* expression is regulated is unclear. In *Bacillus subtilis*, flagellin homeostasis and motility are coregulated by flagellar assembly factor FliW, Flagellin Hag (FliC homolog), and CsrA (Carbon storage regulator A), which is referred to as partner-switching mechanism “FliW-CsrA-Hag”. In this study, we characterized FliW and CsrA functions by deleting or overexpressing *fliW*, csrA, and *fliW*-*csrA* in *C. difficile* R20291. We showed that both *fliW* deletion or *csrA* overexpression in R20291, and *csrA* complementation in R20291ΔWA (*fliW*-*csrA* codeletion) dramatically decreased FliC production, however, *fliC* gene transcription was unaffected. While suppression of *fliC* translation by *csrA* overexpression was mostly relieved when *fliW* was coexpressed, and no significant difference in FliC production was detected when only *fliW* was complemented in R20291ΔWA. Further, loss of *fliW* led to increased biofilm formation, cell adhesion, toxin production, and pathogenicity in a mouse model of *C. difficile* infection (CDI), while *fliW*-*csrA* codeletion decreased toxin production and mortality *in vivo*. Taken together, these data suggest that CsrA negatively modulates *fliC* expression and FliW indirectly affects *fliC* expression through inhibition of CsrA post-transcriptional regulation, which seems similar to the “FliW-CsrA-Hag” switch in *B. subtilis*. Our data also suggest that “FliW-CsrA-*fliC*/FliC” can regulate many facets of *C. difficile* R20291 pathogenicity.

**IMPORTANCE:** *C. difficile* flagellin FliC is associated with toxin gene expression, bacterial colonization and virulence, and is also involved in pleiotropic gene regulation during *in vivo* infection. However, how f*liC* expression is regulated remains unclear. In light of “FliW-CsrA-Hag” switch coregulation mechanism reported in *B. subtilis*, we showed that *fliW* and *csrA* play an important role in flagellin synthesis which affects *C. difficile* motility directly. Our data also suggest that FliW-CsrA-*fliC*/FliC” can regulate many facets of *C. difficile* R20291 pathogenicity. These findings further aid us in understanding the virulence regulation in *C. difficile*.

## INTRODUCTION

*Clostridioides difficile* (formerly *Clostridium difficile*) (1, 2) is a Gram-positive, spore-forming, toxin-producing, anaerobic bacterium that is a leading cause of nosocomial antibiotic-associated diarrhea in the developed countries (3). *C. difficile* infection (CDI) can result in a spectrum of symptoms, ranging from mild diarrhea to pseudomembranous colitis and potential death (4). *C. difficile* has many virulence factors, among which toxin A (TcdA) and toxin B (TcdB) are the major ones (5, 6). These toxins can disrupt the actin cytoskeleton of intestinal cells through glucosylation of the Rho family of GTPases, and induce mucosal inflammation and symptoms associated with CDI (7).

The carbon storage regulator A (CsrA) has been reported to control various physiological processes, such as flagella synthesis, virulence, central carbon metabolism, quorum sensing, motility, biofilm formation in pathogens including *Pseudomonas aeruginosa, Pseudomonas syringae, Borrelia burgdorferi, Salmonella typhimurium*, and *Proteus mirabilis* (8-14). It is a widely distributed RNA binding protein that post-transcriptionally modulates gene expression through regulating mRNA stability and / or translation initiation of target mRNA (13, 15). CsrA typically binds to multiple specific sites that are located nearby or overlapping the cognate Shine-Dalgarno (SD) sequence in the target transcripts (16, 17). The roles of CsrA in *Bacillus subtilis* have been also reported (17-20). Yakhnin et al. (17) first reported that CsrA in *B. subtilis* can regulate translation initiation of the flagellin (hag) by preventing ribosome binding to the *hag* transcript. Meanwhile, two CsrA binding sites (BS1: A51 to A55; BS2: C75 to G82) were identified in the *hag* leader of mRNA, among which BS2 overlaps with the *hag* mRNA SD sequence. Mukherjee et al. (18) elucidated that the interaction between CsrA and FliW could govern flagellin homeostasis and checkpoint on flagellar morphogenesis in *B. subtilis*. FliW, the first protein antagonist of CsrA activity was also identified and characterized in *B. subtilis*. They elegantly demonstrated a novel regulation system “a partner-switching mechanism” (Hag-FliW-CsrA) on flagellin synthesis in *B. subtilis*. Briefly, following the flagellar assembly checkpoint of hook completion, FliW was released from a FliW-Hag complex. Afterward, FliW binds to CsrA which will relieve CsrA-mediated *hag* translation repression for flagellin synthesis concurrent with filament assembly. Thus, flagellin homeostasis restricts its own expression on the translational level. Results also suggested that CsrA has an ancestral role in flagella assembly and has evolved to coregulate multiple cellular processes with motility. Oshiro et al. (19) further quantitated the interactions in the Hag-FliW-CsrA system. They found that Hag-FliW-CsrA^dimer^ functions at nearly 1:1:1 stoichiometry. The Hag-FliW-CsrA^dimer^ system is hypersensitive to the cytoplasmic Hag concentration and is robust to perturbation.

Recently, the role of CsrA on carbon metabolism and virulence associated processes in *C. difficile* 630Δerm was analyzed by overexpressing the *csrA* gene (20). Authors showed that the *csrA* overexpression can increase motility ability, toxin production, and cell adherence, and induce carbon metabolism change. *C. difficile* flagellin gene *fliC* is associated with toxin gene expression, bacterial colonization, and virulence, and is responsible for pleiotropic gene regulation during *in vivo* infection (21-25). The delicate regulations among *fliC* gene expression, toxin production, bacterial motility, colonization, and pathogenicity in *C. difficile* are indicated. Though the important roles of CsrA in flagellin synthesis and flagellin homeostasis have been studied in other bacteria (17-19), the regulation of FliW, CsrA, and FliC and the function of *fliW* in *C. difficile* remain unclear.

In this communication, we aimed to study the involvement of FliW and CsrA in *fliC* expression and *C*.*difficile* virulence and physiology by constructing and analyzing *fliW* and *fliW*-*csrA* deletion mutants of *C. difficile* R20291. We evaluated these mutants in expression of *fliC*, motility, adhesion, biofilm formation, toxin production, sporulation, germination, and pathogenicity in a mouse model of CDI.

## RESULTS

### Construction of *fliW* and *fliW*-*csrA* deletion mutants and complementation strains

The *C. difficile* R20291 flagellar gene operon was analyzed through the *IMG*/*M* website (https://img.jgi.doe.gov/), and the late-stage flagellar genes (F1) were drawn as Fig. 1A (21). Among them, *fliW* and *csrA* genes have a 10 bp overlap and were demonstrated as cotranscription by RT-PCR (Fig. S1).

**Fig. 1.**
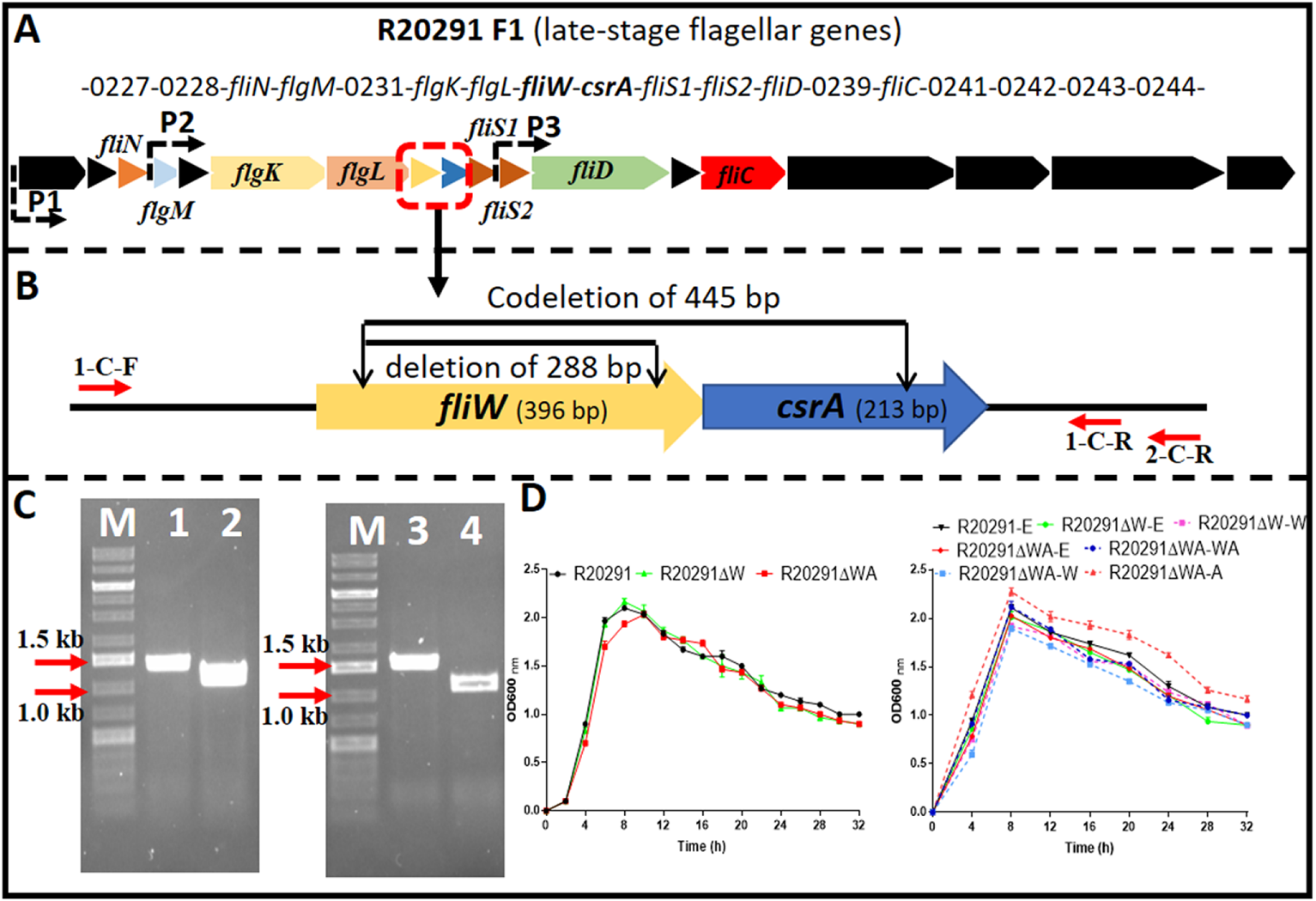
R20291 late-stage flagellar genes (F1) and *fliW* and *fliW*-*csrA* deletions. (A) Schematic representation of late-stage flagellar genes (F1). Dotted arrows (P1, P2, and P3) indicate the potential promoters in F1. (B) Deletion of *fliW* and *fliW*-*csrA* genes. 1-C-F/R were used to verify *fliW* deletion, and 1-C-F and 2-C-R were used to test *fliW*-*csrA* codeletion. (C) Verification of *fliW* and *fliW*-*csrA* deletions by PCR. M: DNA ladder; 1: R20291 genome as PCR template; 2: R20291ΔW genome as PCR template; 3: R20291 genome as PCR template; 4: R20291ΔWA genome as PCR template. (D) Growth profile of parent strain and gene deletion mutants. Experiments were independently repeated thrice. Bars stand for mean ± SEM. One-way ANOVA with post-hoc Tukey test was used for statistical significance.

To analyze the role of *fliW* and *csrA* in R20291, CRISPR-AsCpfI based plasmid pDL1 (pMTL82151-Ptet-AscpfI) was constructed for gene deletion in *C. difficile* (26, 27). pDL1-*fliW* and pDL1-*csrA* gene deletion plasmids were constructed, and the *fliW* gene (288 bp deletion) (R20291Δ*fliW*, referred hereafter as R20291ΔW) was deleted successfully. However, after several trials, we couldn’t get the *csrA* gene deletion mutant possibly due to its small size (213 bp). Therefore, *fliW*-*csrA* codeletion plasmid pDL1-*fliW*-*csrA* was constructed and the *fliW*-*csrA* (445 bp deletion) codeletion mutant (R20291Δ*fliW*-*csrA*, referred hereafter as R20291ΔWA) was obtained (Fig. 1 B and C). To study the role of *csrA* in R20291, the single gene complementation strain R20291ΔWA-W and R20291ΔWA-A were constructed. R20291, R20291-pMTL84153 (R20291-E), R20291ΔW-pMTL84153 (R20291ΔW-E), and R20291ΔWA-pMTL84153 (R20291ΔWA-E) were used as control strains when needed.

The effects of *fliW* and *fliW*-*csrA* deletion on R20291 growth were evaluated. Fig. 1D showed that there was no significant difference in bacterial growth between wild type strain and mutants in BHIS media.

### Effects of *fliW* and *fliW*-*csrA* deletions on *C. difficile* motility and biofilm formation

To characterize the effects of *fliW* and *fliW*-*csrA* deletions on *C. difficile* motility, swimming (Fig. 2A; Fig. S2) and swarming (Fig. S2) motilities of R20291, R20291ΔWA, and R20291ΔW were first analyzed at 24 h and 48 h post-inoculation, respectively. The diameter of the swimming halo of R20291ΔWA increased by 27.2% (*p* < 0.05), while that of R20291ΔW decreased by 58.4% (*p* < 0.05) compared to that of R20291. Next, we examined the motility of the complementation strains (Fig. 2B; Fig. S2), and similar results were obtained among R20291-E, R20291ΔWA-E (with the swimming halo increased by 74.8%, *p* < 0.05), and R20291ΔW-E (with the swimming halo decreased by 59.2%, *p* < 0.05) (Fig. 2B). No significant difference was detected between complementation strain R20291ΔWA-WA, R20291ΔWA-W, R20291ΔW-W, and the parent strain R20291-E except R20291ΔWA-A which decreased by 52.0% (*p* < 0.05) in swimming halo (Fig. 2B). The swarming (48 h) and swimming (24 h) motilities analyzed on agar plates were shown in Fig. S2.

**Fig. 2.**
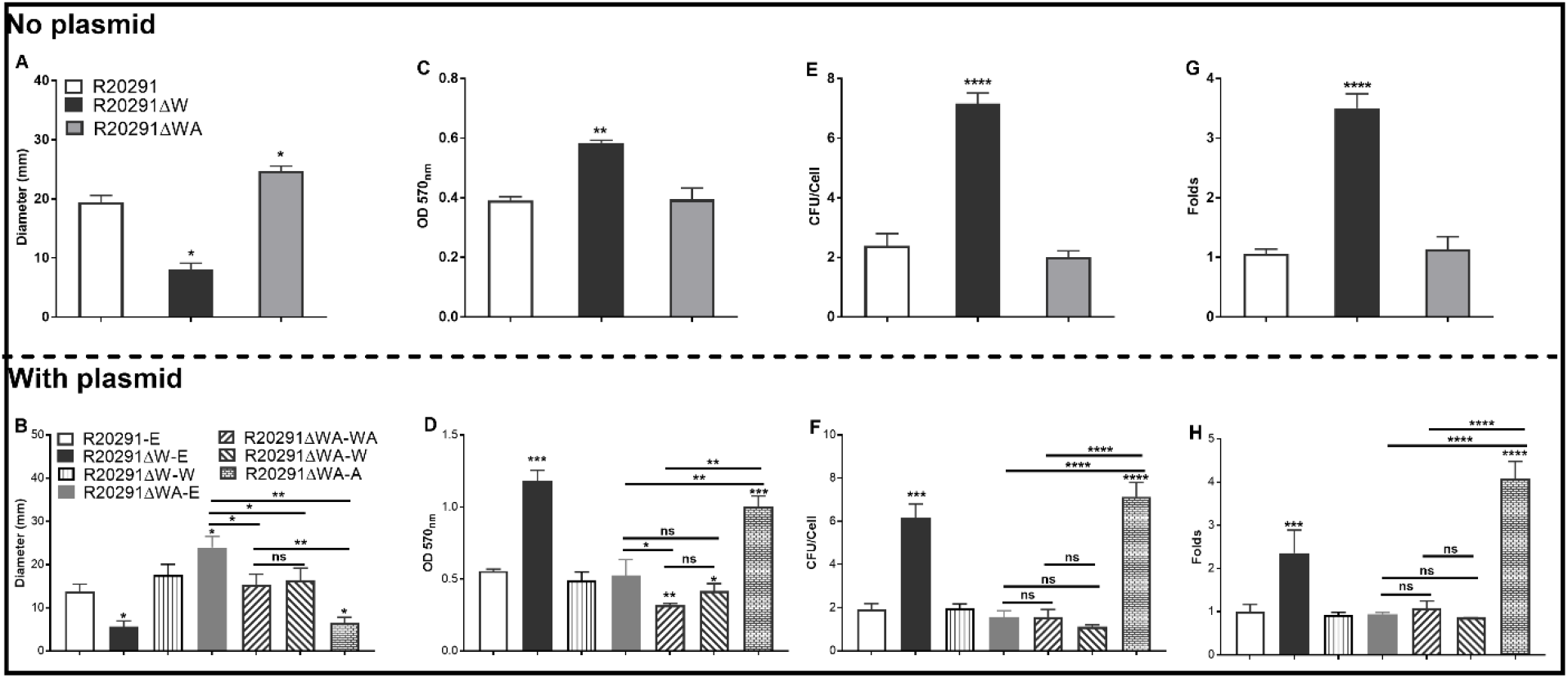
Motility, biofilm, and adhesion analysis. (A) and (B): Halo diameter of motility (swimming analysis on 0.175% agar plate). (C) and (D): Biofilm formation analysis. (E) an (F): Adherence of *C. difficile* vegetative cells to HCT-8 cells *in vitro*. (G) and (H): Adhesion analysis with 5(6)-CFDA dye. The fluorescence intensity was scanned by the Multi-Mode Reader (excitation, 485 nm; emission, 528 nm). The original relative fluorescence unit (RFU) was recorded as F0, after PBS wash, the RFU was recorded as F1. The adhesion ratio was calculated as follows: F1/F0. Experiments were independently repeated thrice. Bars stand for mean ± SEM (\**P* < 0.05, \*\**P* < 0.01, \*\*\**P* < 0.001, \*\*\*\**P* < 0.0001). One-way ANOVA with post-hoc Tukey test was used for statistical significance. * directly upon the column means the significant difference of the experimental strain compared to R20291 or R20291-E.

The effects of *fliW* and *fliW*-*csrA* deletions on *C. difficile* biofilm formation were also analyzed. In comparison with R20291, the biofilm formation of R20291ΔW increased by 49.5% (*p* < 0.01), and no significant difference in biofilm formation was detected in R20291ΔWA (Fig. 2C). The biofilm formation of R20291ΔW-E increased 1.12 fold (*p* < 0.001) and R20291ΔWA-A increased by 79.9% (*p* < 0.001) compared to R20291-E (Fig. 2D). Meanwhile, the biofilm formation of R20291ΔWA-WA and R20291ΔWA-W decreased by 42.8% (*p* < 0.01) and 25.2% (*p* < 0.05), respectively.

Together, these data indicate that loss of FliW impairs *C. difficile* motility, and increases biofilm production. The decrease of motility and increase of biofilm production were also detected in R20291ΔWA-A, which was largely restored by coexpressing *fliW* with *csrA* in R20291ΔWA (Fig. 2B; Fig. 2D), indicating that *fliW* works together with *csrA* to regulate bacterial motility and biofilm production.

### Effects of *fliW* and *fliW*-*csrA* deletions on bacterial adherence *in vitro*

The ability of *C. difficile* vegetative cells to adhere to HCT-8 cells *in vitro* was analyzed. Fig. 2E showed that the mean adhesion number of R20291 was 2.40 ± 0.70 bacteria / cell, while that of R20291ΔW was 7.17 ± 0.61, which was 3.0 fold (*P* < 0.0001) of R20291. No significant difference was detected between R20291ΔWA and R20291. In the complementation strains, we detected a similar result which showed that the mean adhesion number of R20291ΔW-E (6.17 ± 0.64) was 3.20 fold (*P* < 0.0001) of R20291-E (1.93 ± 0.25) (Fig. 2F). The adhesion ability of complementation strains nearly recovered to that of wild type strain except for R20291ΔWA-A (7.13 ± 0.66, *P* < 0.0001) which was 3.69 fold of R20291-E in the mean adhesion number (Fig. 2F).

To visualize the adhesion of *C. difficile* to HCT-8 cells, the *C. difficile* vegetative cells were labeled with the chemical 5(6)-CFDA. Fig. 2G and 2H showed that the fluorescence intensity of R20291ΔW was 3.50 fold (*P* < 0.0001) of that in R20291, and the fluorescence intensity of R20291ΔW-E was 2.36 fold (*P* < 0.001) and R20291ΔWA-A was 4.08 fold (*P* < 0.0001) of that in R20291-E, respectively, which is consistent with the results showed in the Fig. 2E and 2F. Meanwhile, the adherence of *C. difficile* to HCT-8 cells was also visualized by fluorescence microscopy (Fig. S3).

Our data showed that FliW negatively affects bacterial adherence. CsrA complementation in R20291ΔWA increased adherence, while the phenotype change can be recovered partially when *fliW* was coexpressed with *csrA* in R20291ΔWA, suggesting that *fliW* works together with *csrA* to regulate bacterial adherence. The results from bacterial adherence analysis were consistent with biofilm production analysis indicating the close relation between biofilm production and adherence in *C. difficile*.

### Effects of deletion and overexpression of *fliW* and *fliW*-*csrA* on *fliC* expression

In *B. subtilis*, FliW interacts with CsrA to regulate *hag* (a homolog of *fliC*) expression. We reasoned that FliW and CsrA would also regulate *fliC* expression in *C. difficile*. As shown in Fig. 3A, the transcription of *fliC* in R20291ΔWA increased 1.12 fold (*p* < 0.05), while the *fliW* deletion impaired the *fliC* transcription slightly while no significant difference. Fig. 3B showed the production of FliC in R20291ΔW dramatically decreased (10.4 fold reduction, *p* < 0.001), while that of R20291ΔWA increased significantly (increased by 27.5%, *p* < 0.05). To further determine the role of the single gene *csrA* on FliC synthesis, the *csrA* and *fliW* were complemented into R20291ΔWA or overexpressed in R20291, respectively. Results showed that the significant difference of *fliC* transcription could only be detected in R20291ΔWA-E (increased by 32.3%, *p* < 0.05) (Fig. 3C) and R20291-W (increased by 69.8%) compared to R20291-E (Fig. 3E). Interestingly, the FliC production of R20291ΔWA-A was 4.2 fold (*p* < 0.001) of that in R20291-E, while that of R20291ΔWA-WA only decreased by 14.3% (*p* < 0.05) and no significant difference of FliC production in R20291ΔWA-W was detected (Fig. 3D). As shown in Fig. 3E and 3F, the *fliC* transcription of R20291-A was not affected compared to R20291-E, but the FliC production in R20291-A decreased 5.3 fold (*p* < 0.0001). The decrease of FliC production in R20291-A can be partially recovered when *fliW* was coexpressed with *csrA* (R20291-WA decreased by 16.2%, *p* < 0.05).

**Fig. 3.**
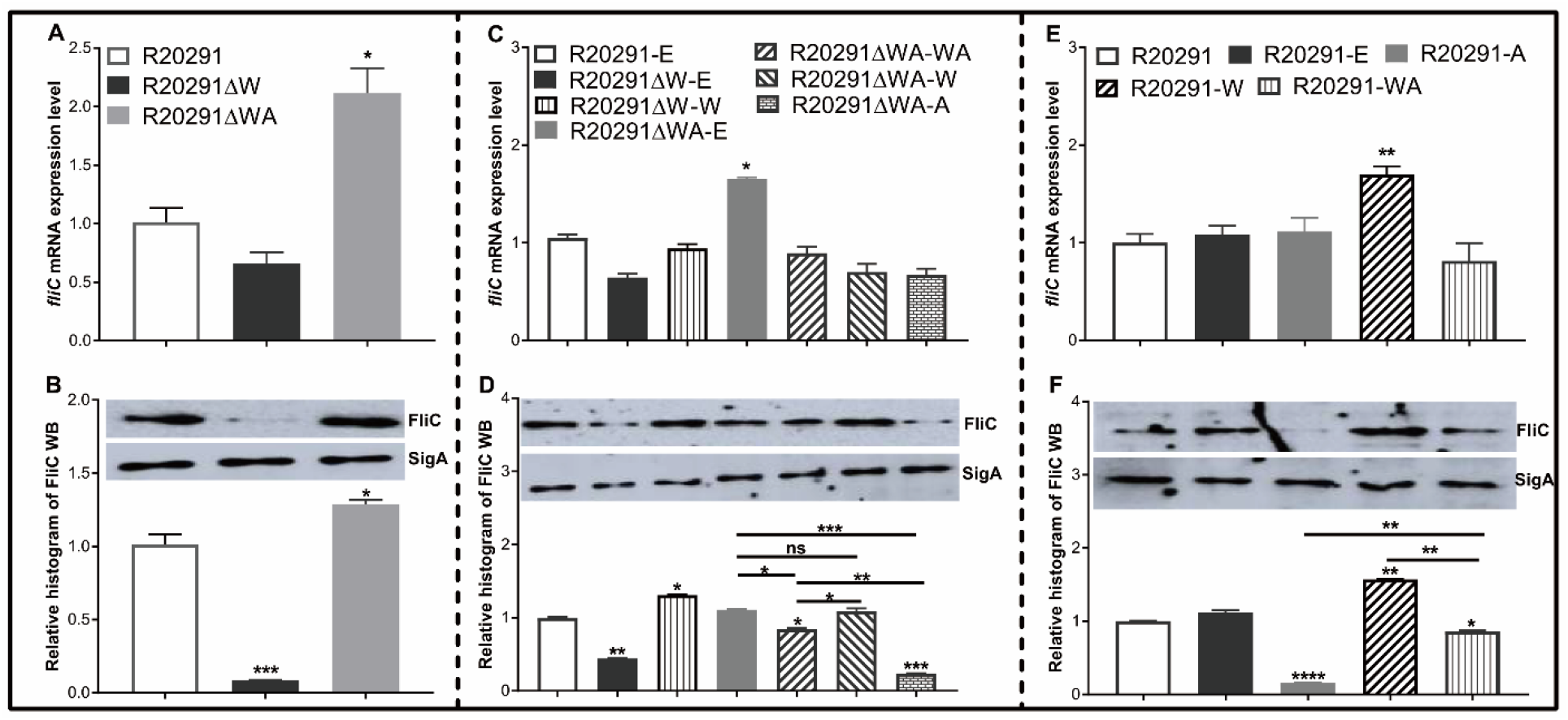
*fliC* expression analysis. (A), (C), and (E) Analysis of *fliC* expression on transcription level. (B), (D), and (F) Analysis of *fliC* expression on translation level by Western blot. SigA protein was used as a loading control. Experiments were independently repeated thrice. Bars stand for mean ± SEM (\**P* < 0.05, \*\**P*<0.01, \*\*\**P* < 0.001, \*\*\*\**P* < 0.0001). One-way ANOVA with post-hoc Tukey test was used for statistical significance. * upon the column directly means the significant difference of experimental strain compared to R20291 or R20291-E.

Collectively, our data indicate that CsrA negatively modulates *fliC* expression post-transcriptionally and FliW works against CsrA to regulate *fliC* expression possibly through inhibiting CsrA-mediated negative post-transcriptional regulation.

### Effects of *fliW* and *fliW*-*csrA* deletions on toxin expression

It has been reported that the expression of *csrA* could affect toxin expression in *C. difficile* (20). To evaluate the effects of *fliW* and *fliW*-*csrA* deletions on toxin production, the supernatants of *C. difficile* cultures were collected at 24 and 48 h post-inoculation, and the toxin concentration was determined by ELISA. Fig. 4A showed that the TcdA concentration of R20291ΔWA decreased by 28.6% (*P* < 0.05), while R20291ΔW increased by 65.1% (*P* < 0.01) compared to R20291 at 24 h post-inoculation. However, after 48 h incubation, no significant difference was detected. In Fig. 4B, TcdB concentration of R20291ΔWA decreased by 26.4% (*P* < 0.05) at 24 h post-inoculation, while that of R20291ΔW increased by 93.6% (*P* < 0.01) at 24 h and 33.0% (*P* < 0.05) at 48 h. Similar results were also detected in the complementation strains group (Fig. 4C and 4D). As shown in Fig. 4C and 4D, after 24 h post-inoculation, TcdA (Fig. 4C) concentration of R20291ΔWA-E and R20291ΔWA-W decreased by 33.0% (\**P* < 0.05) and 47.7% (*P* < 0.01), and TcdB (Fig. 4D) concentration of R20291ΔWA-E and R20291ΔWA-W decreased by 37.9% (*P* < 0.05) and 31.3% (*P* < 0.05), respectively. While TcdA concentration of R20291ΔW-E, R20291ΔWA-A, and R20291ΔW-W increased by 83.1% (*P* < 0.01), 64.7% (*P* < 0.05), and 56.5% (*P* < 0.05), respectively. Meanwhile, TcdB concentration of R20291ΔW-E increased by 100.2% (*P* < 0.01). At 48 h post-inoculation, though no significant difference in TcdA production was detected among different *C. difficile* strains, TcdB concentration of R20291ΔWA-A increased by 28.5% (*P* < 0.05) compared to R20291-E.

**Fig. 4.**
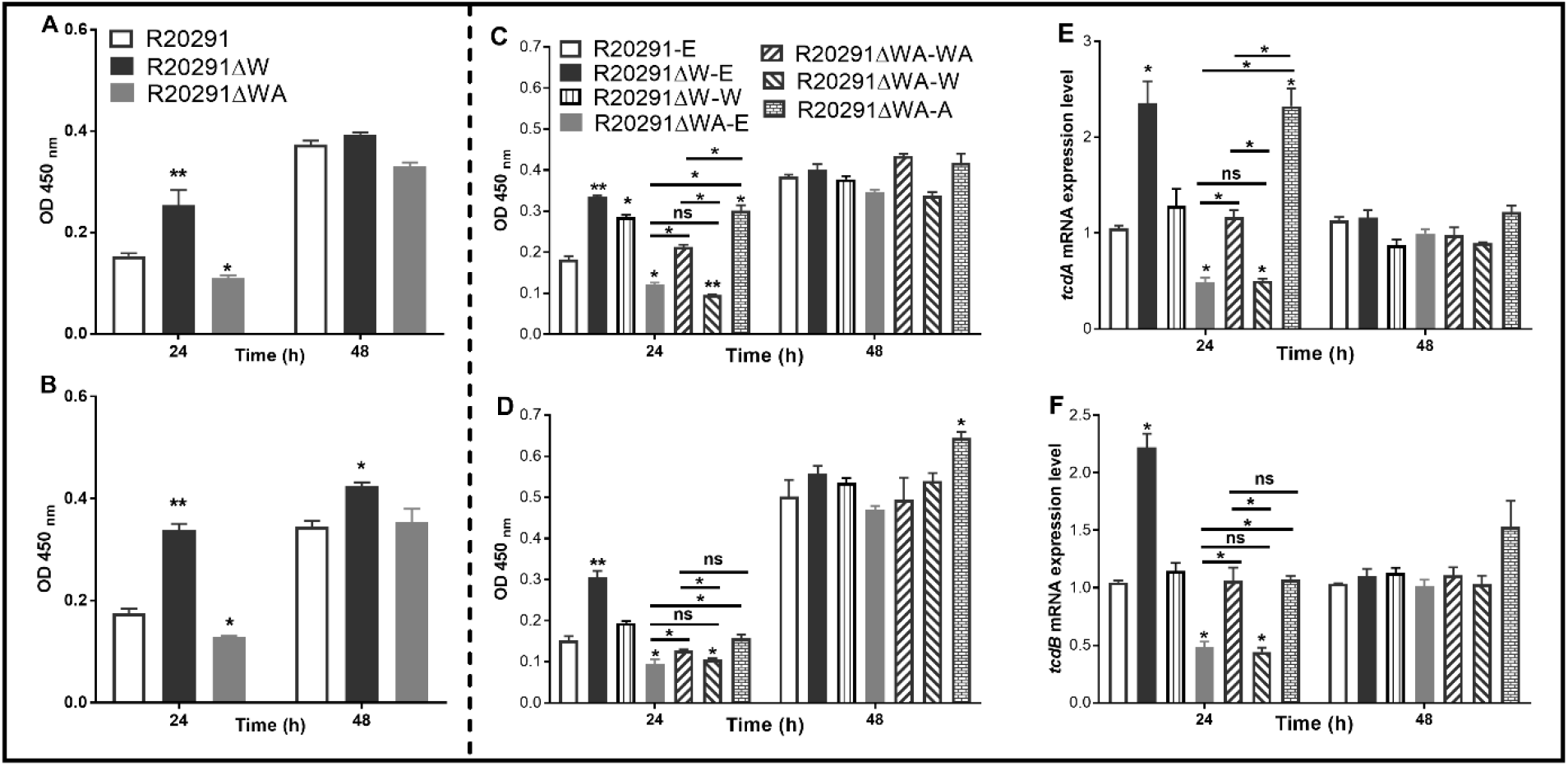
Toxin expression analysis. (A) TcdA concentration in the supernatants of R20291, R20291ΔWA, and R20291ΔW. (B) TcdB concentration in the supernatants of R20291, R20291ΔWA, and R20291ΔW. (C) TcdA concentration in the supernatants of parental and gene complementation strains. (D) TcdB concentration in the supernatants of parental and gene complementation strains. (E) Transcription of *tcdA* in the supernatants of parental and gene complementation strains. (F) Transcription of *tcdB* in the supernatants of parental and gene complementation strains. Experiments were independently repeated thrice. Bars stand for mean ± SEM (\**P* < 0.05, \*\**P* < 0.01). One-way ANOVA with post-hoc Tukey test was used for statistical significance. * upon the column directly means the significant difference of experimental strain compared to R20291 or R20291-E.

To analyze the transcription of *tcdA* and *tcdB* in the complementation strains, RT-qPCR was performed. As shown in Fig. 4E and 4D, the transcription of *tcdA* and *tcdB* of R20291ΔWA-E and R20291ΔWA-W decreased significantly (*P* < 0.05), while that of R20291ΔW-E increased significantly (*P* < 0.05). Interestingly, the *tcdA* transcription of R20291ΔWA-A also showed a significant increase (*P* < 0.05) compared to the wild type strain. Our data indicate that FliW negatively regulates toxin expression, while CsrA plays a positive regulation role in toxin expression.

### Effects of *fliW* and *fliW*-*csrA* deletions on sporulation and germination

To assay the sporulation ratio of *C. difficile* strains, R20291, R20291ΔWA, and R20291ΔW were cultured in Clospore media for 48 and 96 h, respectively. Results (Fig. S4A) showed that no significant difference in the sporulation ratio was detected between the wild type strain and the mutants. The germination ratio of *C. difficile* spores was evaluated as well. Purified spores of R20291, R20291ΔWA, and R20291ΔW were incubated in the germination buffer supplemented with taurocholic acid (TA). As shown in Fig. S4B, there was no significant difference in the germination ratio between the wild type strain and the mutants.

### Evaluation of *fliW* and *fliW*-*csrA* deletions on bacterial virulence in the mouse model of CDI

To evaluate the effects of *fliW* and *fliW*-*csrA* deletions on *C. difficile* virulence *in vivo*, the mouse model of CDI was used. Thirty mice (n=10 per group) were orally challenged with R20291, R20291ΔWA, or R20291ΔW spores (1 × 10^6^ spores / mouse) after antibiotic treatment. As shown in Fig. 5A, the R20291ΔW infection group lost more weight at post challenge days 1 (*P* < 0.05) and the R20291ΔWA infection group lost less weight at post challenge days 3 (*P* < 0.05) compared to the R20291 infection group. Fig. 5B showed that 60% of mice succumbed to severe disease within 4 days in the R20291ΔW infection group and 20% in the R20291ΔWA infection group compared to 50% mortality in the R20291 infection group (no significant difference with log-rank analysis). Meanwhile, 100% of mice developed diarrhea in both the R20291ΔW and R20291 infection groups versus 80% in the R20291ΔWA infection group at post challenge days 2 (Fig. 5C). As shown in Fig. 5D, the CFU of the R20291ΔW infection group increased in the fecal shedding samples at post challenge days 1 and 2 (*P* < 0.05), while the CFU of the R20291ΔWA infection group decreased at post challenge days 1, 5, and 6 (*P* < 0.05) compared to the R20291 infection group.

**Fig. 5.**
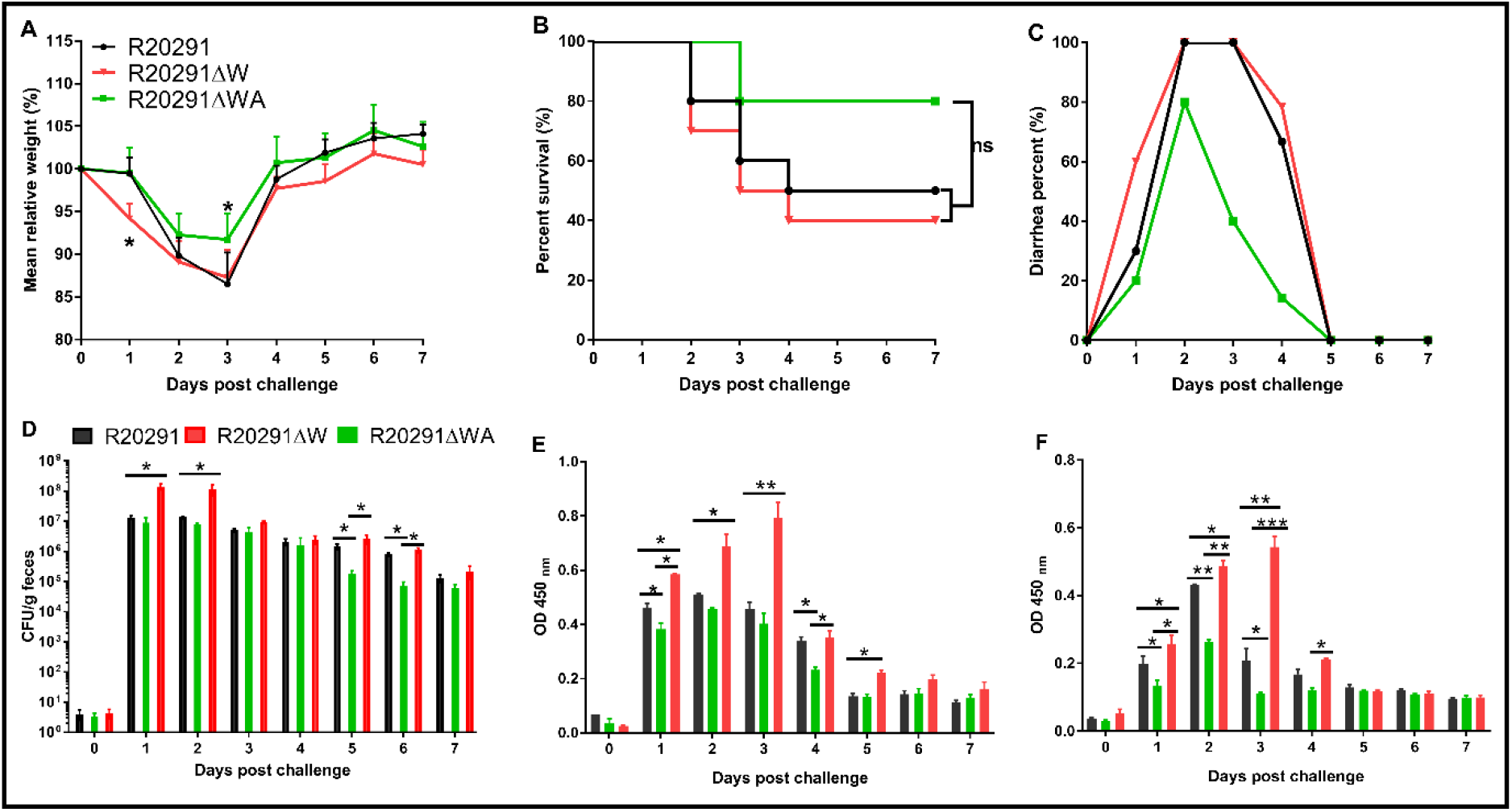
Effects of *fliW* and *fliW-csrA* deletion on *C. difficile* virulence in mice. (A) Mean relative weight changes. (B) Survival curve. (C) Diarrhea percentage. (D) *C. difficile* in feces. (E) TcdA titer of fecal sample. (F) TcdB titer of fecal sample. Bars stand for mean ± SEM (\**P* < 0.05, \*\**P* < 0.01). One-way ANOVA with post-hoc Tukey test was used for statistical significance. Animal survivals were analyzed by Kaplan-Meier survival analysis with a log-rank test of significance.

To evaluate the toxin level in the gut, the concentration of TcdA and TcdB in the feces was measured. In comparison with the R20291 infection group, the TcdA of the R20291ΔW infection group increased significantly at post challenge days 1 (*P* < 0.05), 2 (*P* < 0.05), 3 (*P* < 0.01), and 5 (*P* < 0.05) (Fig. 5E). While the TcdA of the R20291ΔWA infection group was decreased significantly at post challenge days 1 (*P* < 0.05) and 4 (*P* < 0.05) (Fig. 5E). As shown in Fig. 5F, the TcdB concentration of the R20291ΔW infection group decreased significantly at post challenge days 1 (*P* < 0.05), 2 (*P* < 0.05), and 3 (*P* < 0.05), and that of the R20291ΔWA increased significantly at post challenge days 1 (*P* < 0.05), 2 (*P* < 0.01), and 3 (*P* < 0.01). Taken together, our results indicate that the FliW defect increases R20291 pathogenicity *in vivo*, while the *fliW*-*csrA* codeletion impairs R20291 pathogenicity.

## DISCUSSION

In this study, we sought to characterize the impacts of FliW, CsrA, and FliC on *C. difficile* pathogenicity. Our data suggest that CsrA negatively modulates *fliC* expression post-transcriptionally and FliW affects *fliC* expression possibly through inhibiting CsrA-mediated negative post-transcriptional regulation. Our data also indicate that FliW negatively affects *C. difficile* pathogenicity possibly by antagonizing CsrA *in vivo*. Based on our current pleiotropic phenotype analysis, a similar partner-switching mechanism “FliW-CsrA-*fliC*/FliC” is predicted in *C. difficile*, though more direct experimental data are needed to uncover the molecular interactions of CsrA, FliW, and *fliC*/FliC in *C. difficile* (Fig. S5).

It has been reported that overexpression of the *csrA* gene could result in flagella defects, poor motility, and increased toxin production and adhesion in *C. difficile* 630Δerm (20). We found that *fliW* and *csrA* genes are broadly found in the *C. difficile* genomes, among them 10 different *C. difficile* strains from ribotype 106 (RT106), RT027, RT001, RT078, RT009, RT012, RT046, and RT017 were selected and compared to R20291 (Table S2). CsrA and FliW widely exist in *C. difficile*, even in the *C. difficile* strains without flagellar like *C. difficile* M120 (28), indicating a potentially important role of FliW-CsrA in *C. difficile*. Interestingly, while there is no flagellar in *C. difficile* M120, but 6 flagellar structure genes (*fliS, fliN, flgK, flgL, fliC*, and *fliD*) are still found in the genome, which inspired us to explore the potential roles of *fliW, csrA*, and *fliC* in *C. difficile* by deleting or overexpressing *fliW, csrA*, and *fliW*-*csrA* genes. The important roles of CsrA in flagellin synthesis and flagellin homeostasis have been reported (17-20). A previous study had shown that the overexpression of the *csrA* gene can cause a dramatic motility reduction and a significant Hag decrease, suggesting that CsrA represses the Hag expression (17). FliW (the first protein regulator of CsrA activity) deletion abolished the *B. subtilis* swarming and swimming motility and decreased the number of flagella and flagellar length (18, 29). In this study, we obtained similar results that FliW defect impaired R20291 motility significantly (Fig. 2A) and increased biofilm formation (Fig. 2C and 2D). Interestingly, the *csrA* gene complementation in R20291ΔWA dramatically suppressed bacterial motility and showed a similar result to R20291ΔW. Inversely, the *fliW*-*csrA* codeletion increased R20291 motility. Meanwhile, no significant difference was detected between R20291ΔWA-W and R20291ΔWA, but there was a significant change between R20291ΔWA-W and R20291-E, indicating that CsrA can suppress *C. difficile* motility and increase biofilm production, while FliW needs to work together with *csrA* to regulate bacteria motility and biofilm formation.

The partner-switching mechanism “Hag-FliW-CsrA” on flagellin synthesis was elucidated in *B. subtilis* and the intracellular concentration of the flagellar filament protein Hag is restricted tightly by the Hag-FliW-CsrA system (18). To investigate whether FliW and CrsA coregulate the *fliC* expression in *C. difficile*, we evaluated both the transcriptional and translational expression level of *fliC* gene. Our data (Fig. 3) showed that the *fliW* deletion resulted in a 10.4 fold decrease of FliC accumulation, while the *fliW*-*csrA* codeletion increased FliC production, indicating that CsrA could suppress the *fliC* translation and FliW works against CsrA to regulate FliC production. In *csrA, fliW*, and *fliW*-*csrA* overexpression experimental groups, we found that the *csrA* overexpression dramatically decreased FliC production (5.3 fold reduction) and the reduction of FliC production in R20291-A can be partially recovered when *fliW*-*csrA* was coexpressed. The FliW complementation in R20291ΔWA didn’t affect FliC production, but the *fliW* overexpression in R20291 increased FliC production. Taken together, our data suggest that CsrA negatively modulates *fliC* expression post-transcriptionally and FliW works against CsrA to regulate *fliC* expression through inhibiting CsrA-mediated negative post-transcriptional regulation, indicating a similar partner-switching mechanism “FliW-CsrA-FliC” in *C. difficile* (Fig. S5). In *B. subtilis*, two CsrA binding sites (BS1: A51 to A55; BS2: C75 to G82) were identified in the *hag* leader of the mRNA (17). Based on the *hag* 5’ untranslated region (5’-UTR) sequence and CsrA conserved binding sequence, a 91 bp 5’-UTR structure with two potential CsrA binding sites (**BS1:** 5’-TGACAA**GGA**TGT-3’, **BS2**: 5’-CTAA**GGA**GGG**-**3’) of *fliC* gene was predicted (Fig. S6) (30). Recently, it was also reported that cytoplasmic Hag levels play a central role in maintaining proper intracellular architecture, and the Hag-FliW-CsrA^dimer^ system works at nearly 1:1:1 stoichiometry(19). Further studies on the exquisite interactions of CsrA, FliW, and *fliC*/FliC in *C. difficile* are still needed.

Flagella play multiple roles in bacterial motility, colonization, growth, toxin production, and survival optimization (21, 31, 32). Recently, several papers have reported that the flagellar genes can affect toxin expression in *C. difficile*, but results from different research groups were controversial (21-23). It was hypothesized that the regulation of the flagellar genes on toxin expression could be caused by the direct change or loss of flagellar genes (such as *fliC* gene deletion) rather than loss of the functional flagella (21). Future study about *fliC* deletion in M120 will be very interesting and will further address the *fliC* gene function in *C. difficile* as there is no flagellar in RT078 strains. In our study, our data indicate that CsrA negatively modulates *fliC* expression and also plays a positive regulation in toxin expression. Inversely, FliW works against CsrA to regulate *fliC* expression which can negatively regulate toxin production. While studies of flagellar effects on motility and toxin production in *C. difficile* from different groups were controversial, the role of the flagella in *C. difficile* pathogenicity can not be overlooked. Dingle et. al (33) and Baban et. al (23) both showed higher mortality of the *fliC* mutant in the animal model of CDI compared to the wild type strains. Our study showed results similar to the published data suggesting that R20291ΔW whose FilC production was dramatically suppressed exhibited higher fatality, while R20291ΔWA showed a decreased pathogenicity compared to R20291 (Fig. 5). In 2014, Barketi et al. (24) examined the pleiotropic roles of the *fliC* gene in R20291 during colonization in mice. Interestingly, the transcription of *fliW* and *csrA* in the *fliC* mutant was 2.03 fold and 4.36 fold, respectively, of that in R20291 *in vivo* experiment (24), which further corroborated that there is a coregulation among *fliC, fliW*, and *csrA*. Surprisingly, transcription of *treA*, a trehalose-6-phosphate hydrolase, increased 177.63 fold (24). Recently, Collins et al. (34) hypothesized that dietary trehalose can contribute to the virulence of epidemic *C. difficile*. The relationship of FliW, CsrA, FliC, and trehalose metabolization is another interesting question in *C. difficile* and some other carbon metabolism affected by the *fliC* mutation could also facilitate *C. difficile* pathogenesis *in vivo*. Previous studies have also highlighted that the flagella of *C. difficile* play an important role in toxin production, biofilm formation, and bacterial adherence to the host (22, 23, 25, 33, 35). In this study, we showed that the FliW defect led to a significant motility decrease, while the biofilm, adhesion, and toxin production increased significantly. Inversely, R20291ΔWA-W, which can imitate the *csrA* gene deletion, showed an increase in motility and a decrease in biofilm formation, toxin production, and adhesion (Fig. 2, Fig. S2, and Fig. S3).

In conclusion, we characterized the function of FliW and CsrA and showed the pleiotropic functions of FliW and CsrA in R20291. Our data suggest that *fliW* and *csrA* play important roles in flagellin (FliC) synthesis which could contribute to *C. difficile* pathogenicity. Currently, *in vitro* study of the interactions of CsrA, FliW, and *fliC*/FliC in *C. difficile* is underway in our group.

## EXPERIMENTAL PROCEDURES

### Bacteria, plasmids, and culture conditions

Table 1 lists the strains and plasmids used in this study. *C. difficile* strains were cultured in BHIS media (brain heart infusion broth supplemented with 0.5% yeast extract and 0.1% L-cysteine, and 1.5% agar for agar plates) at 37 °C in an anaerobic chamber (90% N_2_, 5% H_2_, 5% CO_2_). For spores preparation, *C. difficile* strains were cultured in Clospore media and purified as described earlier (36). *Escherichia coli* DH5α and *E. coli* HB101/pRK24 were grown aerobically at 37 °C in LB media (1% tryptone, 0.5% yeast extract, 1% NaCl). *E. coli* DH5α was used as a cloning host and *E. coli* HB101/pRK24 was used as a conjugation donor host. Antibiotics were added when needed: for *E. coli*, 15 μg/ml chloramphenicol; for *C. difficile*, 15 μg/ml thiamphenicol, 250 μg/ml D-cycloserine, 50 μg/ml kanamycin, 8 μg/ml cefoxitin, and 500 ng/ ml anhydrotetracycline.

**Table 1.**
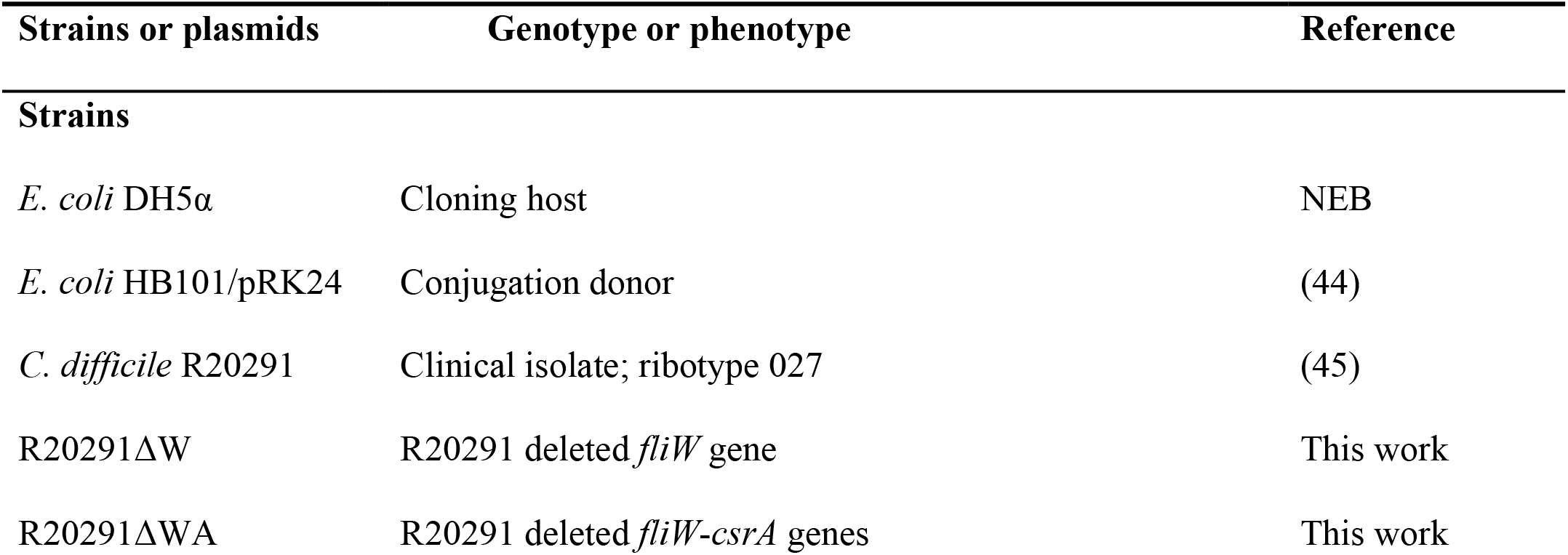
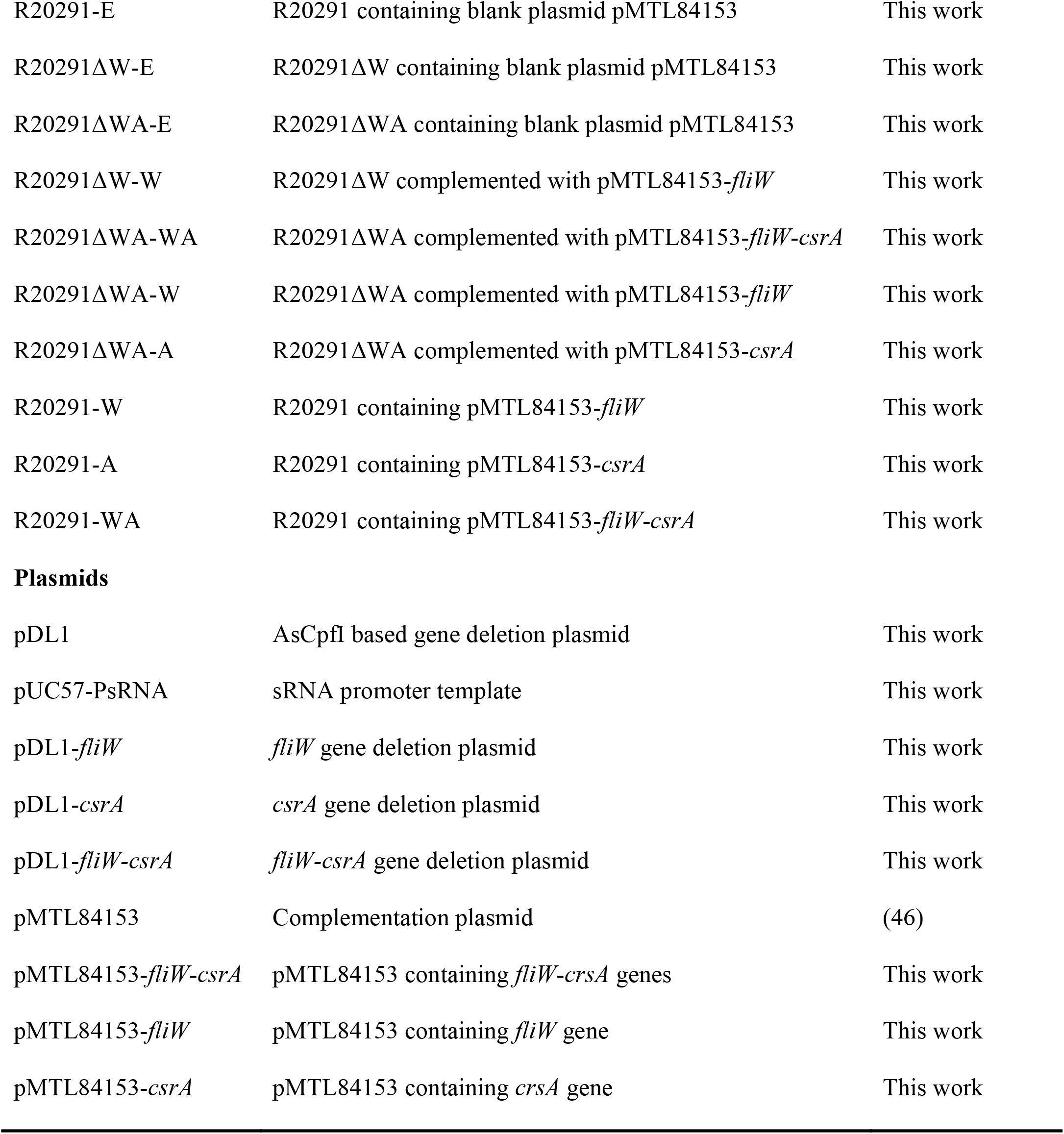
Bacteria and plasmids utilized in this study.

| Strains or plasmids | Genotype or phenotype | Reference |
| --- | --- | --- |
| **Strains** |  |  |
| *E. coli* DH5α | Cloning host | NEB |
| *E. coli* HB101/pRK24 | Conjugation donor | (44) |
| *C. difficile* R20291 | Clinical isolate; ribotype 027 | (45) |
| R20291ΔW | R20291 deleted *fliW* gene | This work |
| R20291ΔWA | R20291 deleted *fliW-csrA* genes | This work |
| R20291-E | R20291 containing blank plasmid pMTL84153 | This work |
| R20291ΔW-E | R20291ΔW containing blank plasmid pMTL84153 | This work |
| R20291ΔWA-E | R20291ΔWA containing blank plasmid pMTL84153 | This work |
| R20291ΔW-W | R20291ΔW complemented with pMTL84153-*fliW* | This work |
| R20291ΔWA-WA | R20291ΔWA complemented with pMTL84153-*fliW-csrA* | This work |
| R20291ΔWA-W | R20291ΔWA complemented with pMTL84153-*fliW* | This work |
| R20291ΔWA-A | R20291ΔWA complemented with pMTL84153-*csrA* | This work |
| R20291-W | R20291 containing pMTL84153-*fliW* | This work |
| R20291-A | R20291 containing pMTL84153-*csrA* | This work |
| R20291-WA | R20291 containing pMTL84153-*fliW-csrA* | This work |
| Plasmids |  |  |
| pDL1 | AsCpfI based gene deletion plasmid | This work |
| pUC57-PsRNA | sRNA promoter template | This work |
| pDL1-*fliW* | *fliW* gene deletion plasmid | This work |
| pDL1-*csrA* | *csrA* gene deletion plasmid | This work |
| pDL1-*fliW-csrA* | *fliW-csrA* gene deletion plasmid | This work |
| pMTL84153 | Complementation plasmid | (46) |
| pMTL84153-*fliW-csrA* | pMTL84153 containing *fliW-csrA* genes | This work |
| pMTL84153-*fliW* | pMTL84153 containing *fliW* gene | This work |
| pMTL84153-*csrA* | pMTL84153 containing *csrA* gene | This work |

### DNA manipulations and chemicals

DNA manipulations were carried out according to standard techniques (37). Plasmids were conjugated into *C. difficile* as described earlier (38). The DNA markers, protein markers, PCR product purification kit, DNA gel extraction kit, restriction enzymes, cDNA synthesis kit, and SYBR Green RT-qPCR kit were purchased from Thermo Fisher Scientific (Waltham, USA). PCRs were performed with the high-fidelity DNA polymerase NEB Q5 Master Mix, and PCR products were assembled into target plasmids with NEBuilder HIFI DNA Assembly Master Mix (New England, UK). Primers (Supporting Information Table S1) were purchased from IDT (Coralville, USA). All chemicals were purchased from Sigma (St. Louis, USA) unless those stated otherwise.

### Construction of R20291 mutant strains of gene deletion, complementation, and overexpression

The Cas12a (AsCpfI) based gene deletion plasmid pDL-1 was constructed and used for *C. difficile* gene deletion (26). The target sgRNA was designed with an available website tool (http://big.hanyang.ac.kr/cindel/) and the off-target prediction was analyzed on the Cas-OFFinder website (http://www.rgenome.net/cas-offinder/). The sgRNA, up and down homologous arms were assembled into pDL-1. Two target sgRNAs for one gene deletion were selected and used for gene deletion plasmid construction in *C. difficile*, respectively. Briefly, the gene deletion plasmid was constructed in the cloning host *E. coli* DH5α and was transformed into the donor host *E. coli* HB101/pRK24, and subsequently was conjugated into R20291. Potential successful transconjugants were selected with selective antibiotic BHIS-TKC plates (15 μg/ml thiamphenicol, 50 μg/ml kanamycin, 8 μg/ml cefoxitin). The transconjugants were cultured in BHIS-Tm broth (15 μg/ml thiamphenicol) to log phase, then the subsequent cultures were plated on the inducing plates (BHIS-Tm-ATc: 15 μg/ml thiamphenicol and 500 ng/ml anhydrotetracycline). After 24 - 48 h of incubation, 20 - 40 colonies were used as templates for colony PCR test with check primers for correct gene deletion colony isolation. The correct gene deletion colony was sub-cultured into BHIS broth without antibiotics and was passaged several times to cure the deletion plasmid, then the cultures were plated on BHIS plates and subsequent colonies were replica plated on BHIS-Tm plates to isolate pure clean gene deletion mutants (R20291ΔW and R20291ΔWA). The genome of R20291ΔW and R20291ΔWA were isolated and used as templates for the PCR test with check primers, and the PCR products were sequenced to confirm the correct gene deletion.

The *fliW* (396 bp) (primers 3-F/R), *csrA* (213 bp) (primers 4-F/R), and *fliW*-*csrA* (599 bp) (primers 5-F/R) genes were amplified and assembled into *Sac*I-*Bam*HI digested pMTL84153 plasmid, yielding the complementation plasmid pMTL84153-*fliW*, pMTL84153-*csrA*, and pMTL84153-*fliW*-*csrA*, and were subsequently conjugated into R20291ΔWA, R20291ΔW, and R20291 yielding complemeantation strain R20291ΔWA/pMTL84153-*fliW* (referred as R20291ΔWA-W), R20291ΔWA/pMTL84153-*csrA* (R20291ΔWA-A), R20291ΔWA/pMTL84153-*fliW*-*csrA* (R20291ΔWA-WA), R20291ΔW/pMTL84153-*fliW* (R20291ΔW-W) and overexpression strain R20291/pMTL84153-*fliW* (R20291-W), R20291/pMTL84153-*csrA* (R20291-A), R20291/pMTL84153-*fliW*-*csrA* (R20291-WA).

### Growth profile, motility, and biofilm assay

*C. difficile* strains were incubated to an optical density of OD_600_ of 0.8 in BHIS media and were diluted to an OD_600_ of 0.2. Then, 1% of the culture was inoculated into fresh BHIS, followed by measuring OD_600_ for 32 h.

To examine the effect of *fliW* and *fliW*-*csrA* deletion on *C. difficile* motility, R20291, R20291ΔWA, and R20291ΔW were cultured to an OD_600_ of 0.8. For swimming analysis, 2 µl of *C. difficile* culture was penetrated into soft BHIS agar (0.175%) plates, meanwhile, 2 µl of culture was dropped onto 0.3% BHIS agar plates for swarming analysis. The swimming assay plates were incubated for 24 h and the swarming plates were incubated for 48 h, respectively.

For biofilm formation analysis, wild type and mutant *C. difficile* R20291 strains were cultured to an OD_600_ of 0.8, and 1% of *C. difficile* cultures were inoculated into Reinforced Clostridial Medium (RCM) with 8 well repeats in a 96-well plate and incubated in the anaerobic chamber at 37 °C for 48 h. Biofilm formation was analyzed by crystal violet dye. Briefly, *C. difficile* cultures were removed by pipette carefully. Then 100 µl of 2.5% glutaraldehyde was added into the well to fix the bottom biofilm, and the plate was kept at room temperature for 30 min. Next, the wells were washed with PBS 3 times and dyed with 0.25% (w/v) crystal violet for 10 min. The crystal violet solution was removed, and the wells were washed 5 times with PBS, followed by the addition of acetone into wells to dissolve the crystal violet of the cells. The dissolved solution was further diluted with ethanol 2 - 4 times and biomass was determined at OD_570_.

### Adherence of *C. difficile* vegetative cells to HCT-8 cells

*C. difficile* adhesion ability was evaluated with HCT-8 cells (ATCC CCL-244) (39). Briefly, HCT-8 cells were grown to 95% confluence (2×10^5^/well) in a 24-well plate and then moved into the anaerobic chamber, followed by infecting with 6 × 10^6^ of log phase of *C. difficile* vegetative cells at a multiplicity of infection (MOI) of 30:1. The plate was cultured at 37 °C for 30 min. After incubation, the infected cells were washed with 300 µl of PBS 3 times, and then suspended in RPMI media with trypsin and plated on BHIS agar plates to enumerate the adhered *C. difficile* cells. The adhesion ability of *C. difficile* to HCT-8 cells was calculated as follows: CFU of adhered bacteria / Total cell numbers.

To visualize the adherence of *C. difficile* to HCT-8 cells, *C. difficile* vegetative cells were labeled with the chemical 5(6)-CFDA (5-(and -6)-Carboxyfluorescein diacetate) (40). Briefly, *C. difficile* strains were cultured to an OD_600_ of 0.8, then washed with PBS 3 times and resuspended in fresh BHIS supplemented with 50 mM 5(6)-CFDA, followed by incubation at 37 °C for 30 min in the anaerobic chamber. After post-incubation, the labeled *C. difficile* cells were collected and washed with PBS 3 times, and then resuspended in RPMI medium. Afterward, the labeled *C. difficile* cells were used for the infection experiment as described above. After 30 min post-infection, the fluorescence of each well was scanned by the Multi-Mode Reader (excitation, 485 nm; emission, 528 nm), the relative fluorescence unit (RFU) was recorded as F0. Following, the plates were washed with PBS 3 times to remove unbound *C. difficile* cells, then the plates were scanned and the RFU was recorded as F1. The adhesion ratio was calculated as follows: F1/F0. After scanning, the infected cell plates were further detected by the fluorescence microscope.

### *fliC* expression assay

For *fliC* transcription analysis, 2 ml of 24 h post inoculated *C. difficile* cultures were centrifuged at 4 °C, 12000×g for 5 min, respectively. Then, the total RNA of different strains was extracted with TRIzol reagent. The transcription of *fliC* was measured by RT-qPCR with primers Q-*fliC*-F/R. All RT-qPCRs were repeated in triplicate, independently. Data were analyzed by the comparative CT (2^-ΔΔCT^) method with 16s rRNA as a control.

To analyze the FliC protein level, *C. difficile* cell lysates from overnight cultures were used for Western blot analysis. Briefly, overnight *C. difficile* cultures were collected and washed 3 times with PBS and then resuspended in 5 ml of distilled water. The suspensions were lysed with TissueLyser LT (Qiagen), followed centrifuged at 4°C, 25000×*g* for 1h. The final pellets were resuspended in 30 μl of PBS and the total protein concentration was measured by using a BCA protein assay (Thermo Scientific, Suwanee, GA). Protein extracts were subjected to 10% SDS-PAGE. Sigma A protein (SigA) was used as a loading control protein in SDS-PAGE. FliC and SigA proteins on the gel were detected with anti-FliC and anti-SigA primary antibody (1:1000) and horseradish peroxidase-conjugated secondary antibody goat anti-mouse (Cat: ab97023, IgG, 1:3000, Abcam, Cambridge, MA) by Western blot, respectively. Anti-FliC antibody used in the Western blot analysis is an anti-FliCD serum, generated in the lab. FliCD is a fusion protein containing *C*.*difficile* FliC and FliD (41).

### Toxin expression assay

To evaluate toxin expression in *C. difficile* strains, 10 ml of *C. difficile* cultures were collected at 24 and 48 h post incubation. The cultures were adjusted to the same density with fresh BHIS. Then the collected *C. difficile* cultures were centrifuged at 4 °C, 8000×*g* for 15 min, filtered with 0.22 μm filters, and used for ELISA. Anti-TcdA (PCG4.1, Novus Biologicals, USA) and anti-TcdB (AI, Gene Tex, USA) were used as coating antibodies for ELISA, and HRP-Chicken anti-TcdA and HRP-Chicken anti-TcdB (Gallus Immunotech, USA) were used as detection antibodies.

For toxin transcription analysis, 2 ml of 24 and 48 h post inoculated *C. difficile* cultures were centrifuged at 4 °C, 12000×*g* for 5 min, respectively. Next, the total RNA of different strains was extracted with TRIzol reagent. The transcription of *tcdA* and *tcdB* was measured by RT-qPCR with primers Q-*tcdA*-F/R and Q-*tcdB*-F/R, respectively. All RT-qPCRs were repeated in triplicate, independently. Data were analyzed by using the comparative CT (2^-ΔΔCT^) method with 16s rRNA as a control.

### Germination and sporulation assay

*C. difficile* germination and sporulation analysis were conducted as reported earlier (42). Briefly, for *C. difficile* sporulation analysis, *C. difficile* strains were cultured in Clospore media for 4 days. Afterward, the CFU of cultures from 48 and 96 h were counted on BHIS plates with 0.1% TA to detect sporulation ratio, respectively. The sporulation ratio was calculated as CFU (65 °C heated) / CFU (no heated). For *C. difficile* germination analysis, *C. difficile* spores were collected from 2-week Clospore media cultured bacteria and purified with sucrose gradient layer (50%, 45%, 35%, 25%, 10%). The heated purified spores were diluted to an OD_600_ of 1.0 in the germination buffer [10 mM Tris (pH 7.5), 150 mM NaCl, 100 mM glycine, 10 mM taurocholic acid (TA)] to detect the germination ratio. The value of OD_600_ was monitored immediately (0 min, t_0_), and was detected once every 2 min (t_x_) for 20 min at 37 °C. The germination ratio was calculated as OD_600_ (tx) / OD_600_ (T_0_). Spores in germination buffer without TA were used as the negative control.

### Evaluation of R20291, R20291ΔWA, and R20291ΔW virulence in the mouse model of *C. difficile* infection

C57BL/6 female mice (6 weeks old) were ordered from Charles River Laboratories, Cambridge, MA. All studies were approved by the Institutional Animal Care and Use Committee of University of South Florida. The experimental design and antibiotic administration were conducted as described earlier (43). Briefly, 30 mice were divided into 3 groups in 6 cages. Group 1 mice were challenged with R20291 spores, group 2 mice with R20291ΔWA spores, and group 3 mice with R20291ΔW spores, respectively. Mice were given an orally administered antibiotic cocktail (kanamycin 0.4 mg/ml, gentamicin 0.035 mg/ml, colistin 0.042 mg/ml, metronidazole 0.215 mg/ml, and vancomycin 0.045 mg/ml) in drinking water for 4 days. After 4 days of antibiotic treatment, all mice were given autoclaved water for 2 days, followed by one dose of clindamycin (10 mg/kg, intraperitoneal route) 24 h before spores challenge (Day 0). After that, mice were orally gavaged with 10^6^ of spores and monitored daily for a week for changes in weight, diarrhea, and mortality.

### Evaluation of *C. difficile* spores and determination of toxin level in feces

Fecal pellets from post infection day 0 to day 7 were collected and stored at -80 °C. To enumerate *C. difficile* numbers, feces were diluted with PBS at a final concentration of 0.1 g/ml, followed by adding 900 µl of absolute ethanol into 100 µl of the fecal solution, and kept at room temperature for 1 h to inactivate vegetative cells. Afterward, 200 µl of vegetative cells inactivated fecal solution from the same group and the same day was mixed. Then, fecal samples were serially diluted and plated on BHIS-CCT plates (250 μg/ml D-cycloserine, 8 μg/ml cefoxitin, 0.1% TA). After 48 h incubation, colonies were counted and expressed as CFU/g feces. To evaluate toxin tilter in feces, 0.1 g/ml of the fecal solution was diluted two times with PBS, followed by examining TcdA and TCdB ELISA.

### Statistical analysis

The reported experiments were conducted in independent biological triplicates except for the animal experiment, and each sample was additionally taken in technical triplicates. Animal survivals were analyzed by Kaplan-Meier survival analysis and compared by the Log-Rank test. One-way analysis of variance (ANOVA) with post-hoc Tukey test was used for more than two groups comparison. Results were expressed as mean ± standard error of the mean. Differences were considered statistically significant if *P* < 0.05 (*).

## ACKNOWLEDGEMENTS

This work was supported in part by the National Institutes of Health grants (R01-AI132711 and R01-AI149852). Authors thank Dr. Abhraham L. Sonnenshein at Tufts University, Dr. Joseph Sorg at Texas A &M, and Dr. Daniel Kearns at Indiana University for the gifts *C. difficile* R20291, *E*.*coli* HB101/pRK24, and anti-SigA primary antibody, respectively. We thank Dr. Nigel Minton at the University of Nottingham for the gift plasmids pMTL84151 and pMTL83353. We also thank Jessica Bullock and Dr. Heather Danhof for their mindful revision and comments.

